# Isolation and characterization of novel temperate virus *Aeropyrum* globular virus 1 infecting hyperthermophilic archaeon *Aeropyrum*

**DOI:** 10.1101/2021.04.21.440749

**Authors:** Maho Yumiya, Yuto Fukuyama, Yoshihiko Sako, Takashi Yoshida

## Abstract

We isolate a novel archaeal temperate virus named *Aeropyrum* globular virus 1 (AGV1) from the host *Aeropyrum* culture. Reproduction of AGV1 was induced by adding 20 mM tris-acetate buffer to exponentially growing host cells. Negatively stained virions showed spherical morphology (60 ±2 nm in diameter) similar to *Globuloviridae* viruses. The double-stranded circular DNA genome of AGV1 contains 18,222 bp encoding 34 open-reading frames. No ORFs showed significant similarity with *Globuloviridae* viruses. AGV1 shares three genes, including an integrase gene, with reported spindle-shaped temperate viruses. However we couldn’t detect its integration site in the host genome. Moreover AGV1 seemed not to replicate autonomously because there are no origin recognition boxes in the genome. qPCR results showed that the genome copy number of AGV1 was lower than that of the host genome (10^−3^ copies per host genome). Upon the addition of tris-acetate buffer, a steep increase in the AGV1 genome copy number (9.5–26 copies per host genome at 2 days post-treatment) was observed although clustered regularly interspaced short palindromic repeat (CRISPR) elements of the host genome showed significant matches with AGV1 protospacers. Our findings suggest that AGV1 is a novel globular virus exhibiting an unstable carrier state in the growing host and in that way AGV1 can escape from the host defense system and propagate under stressful host conditions.

**Importance:** Studying archaeal viruses yields novel insights into the roles of virospheres and viruses in the evolutionary process of their hosts. Here, we isolated a novel spherical virus named *Aeropyrum* globular virus 1. AGV1 has integrase gene but its genome is not integrated into the host genome. AGV1 could not replicate autonomously due to the lack of origin recognition boxes and thus its copy number was too low (10^−3^ copies per host genome) without any inducing stimulus. However, upon the addition of tris-acetate buffer, the AGV1 genome copy number steeply increased instead of a perfect sequence match between the spacer of the host CRISPR/Cas system and the protospacer. Our findings suggest that AGV1 can escape from the host defense system and propagate under stressful conditions for the host by establishing an unstable carrier state. These results reveals a novel aspect of host–virus interactions in extreme environments.

## INTRODUCTION

Viruses are absolute intracellular parasites that depend on their host living organisms for replication(1, 2). Through infection, viruses can introduce genetic variation to the host microorganisms (3), affect their host microbial metabolism (4), directly kill their hosts by cell lysis, and accordingly contribute to the diversification of the microbial community (5). Virus– microorganism interactions also drive antagonistic coevolution that increases genetic diversity of both hosts and viruses (2). Thus, viruses are key players in microbial ecology (6).

Most *Crenarchaeota* members are hyperthermophiles that grow optimally at temperatures of at least 80°C. According to the International Committee on Taxonomy of Viruses, to date, cultured crenarchaeal viruses have been identified in 13 genera, such as *Sulfolobus, Acidianus, Stygiolobus, Thermoproteus, Pyrobaculum*, and *Aeropyrum* (7). Crenarchaeal viruses exhibit diverse morphologies, including spindle-shaped and bottle-shaped cells, which have never been reported in other domains of life (7-9). The life cycles of viruses are largely divided into lytic and lysogenic cycles (10). However, to adapt to the harsh conditions of the host habitat, crenarchaeal viruses, except for lytic viruses, e.g., *Thermoproteus tenax* virus 1 (11), *Sulfolobus* turreted icosahedral virus (12) and *Acidianus* two-tailed virus (13), do not exhibit typical lytic cycles and develop a carrier state in their host. Viruses in carrier state represent a harmonious coexistence with host species and propagate without genome integration and cell lysis (8). It is considered that carrier state is unique and significant strategy for viruses infecting *Crenarchaeota*. All members of *Fuselloviridae* (14), *Guttaviridae* (15, 16), and *Bicaudaviridae* (13) are lysogenic viruses and possess integrase genes. Some of them can not only integrate their circular genome into host chromosomes (17) but also establish carrier state. The archaeon *Sulfolobus shibatae* B12 is the natural host of the fusellovirus *Sulfolobus* spindle-shaped virus 1 (SSV1), which exists in episomic and integrated forms. SSV1 can propagate at low levels (10^7^ virus particles/mL) without cell lysis and even in the absence of an inducing stimulus (18). Thus, its lysogenic cycle is a model of carrier state (19). Transcriptomic analysis of SSV1-infected cells suggests that the virus tightly represses gene expression during its carrier state to establish an equilibrium between viral replication and cellular multiplication (20). However, owing to the limited number of cultured viruses and unique genomic contents, knowledge on the mechanisms and the ecological effects of the carrier state is limited.

Belonging to the phylum *Crenarchaeota, Aeropyrum* spp. are aerobic, neutrophilic, and heterotrophic hyperthermophiles (21). There are only two closely related *Aeropyrum* species, *Aeropyrum pernix* (21) and *Aeropyrum camini* (22). Four viruses that infect *A. pernix* have been isolated and described: *Aeropyrum pernix* bacilliform virus 1 (APBV1) (23), *Aeropyrum pernix* ovoid virus 1 (APOV1) (15), *Aeropyrum pernix* spindle-shaped virus 1 (APSV1) (15), and *Aeropyrum* coil-shaped virus (ACV) (24). APBV1 and ACV are the first members of Clavaviridae and Spiraviridae, respectively (23, 24). Further diversity in *A. pernix* viruses was predicted to have been still veiled in hot environments by transmission electron micrograph (TEM) observations (e.g., filamentous virus and short bacilliform virus) (23). *Aeropyrum* species are specialists in their habitat requirements and possess small and very conservative genomes (25). Genomic variation is observed in virus-related elements including two proviral regions (*Fuselloviridae* APSV1 and *Guttaviridae* APOV1), adaptive immune system against foreign genetic elements (clustered regularly interspaced short palindromic repeat [CRISPR]), and ORFans probably originating from viruses (25). In addition, most spacer sequences (141/144) in the CRISPR loci of *A. pernix* K1 and *A. camini* SY1 showed no similarity to databases (25). It is considered that their genomic diversification is mostly derived from viruses and that there are unknown viruses interacting with *A. pernix* and contribute to their population dynamics in the environment.

Here, we report the isolation of a novel temperate virus infecting *Aeropyum* species and investigate how the virus establish its unique carrier state in the host culture using quantitative PCR assay.

## MATERIALS AND METHODS

### Sample collection

On December 12, 2012 and May 27, 2014, samples were collected from the Yamagawa coastal hydrothermal field (31°10’58″ ‘‘ N, 130°36’59’’″ E) in the Kagoshima Prefecture. Effluent seawater and coastal sand were collected using a ladle and stored in 50 mL centrifuge tubes (Greiner Bio-One, frickenhausen,Germany). All tubes were transported in ice packs to the laboratory by a refrigerated courier service (Yamato Transport, Tokyo, Japan) and stored at 4°C until use.

### Isolation of the host strain

To establish the enrichment culture of *A. pernix*, approximately 0.5 g sample collected from 2012 was inoculated into the 5 mL JXTm medium (1 g tryptone, 1 g yeast extract, 28.15 g NaCl, 0.67 g KCl, 5.51 g MgCl_2_·6H_2_O, 6.92 g MgSO_4_·7H_2_O, 1.45 g CaCl_2_, and 1 g Na_2_S_2_O_3_·5H_2_O per liter, pH 7.0) (26) in 18 × 180 mm hermetically sealed screw-cap test tubes. The cultures were incubated at 90°C in a dry oven (FC612 or DRS620DB; ADVANTEC, Tokyo, Japan) under atmospheric conditions. Then, the enriched cells were streaked onto a ST Gelrite plate (containing 32 g of sea salt (Sigma Aldrich, St. Louis, Missouri, USA), 1 g Na_2_S_2_O_3_ 5H_2_O, 0.8 g yeast extract (Becton, Dickinson and Company, Franklin Lakes, New Jersey), 1.2 g tryptone (Becton, Dickinson and Company, Franklin Lakes, New Jersey, USA), and 8 g Gelrite gellan gum (Sigma Aldrich, St. Louis, Missouri, USA) per liter) in glass Petri dishes (10 cm in diameter). The plates were incubated at 90°C in a BBL GasPak™ 100 Holding Jar (Becton, Dickinson and Company, Franklin Lakes, New Jersey, USA) to avoid evaporation. After 3 to 5 days of incubation, well-isolated colonies formed on the surface of the plates were collected and transferred to a fresh JXTm medium. To ensure the purity of the *Aeropyrum* isolate, the streaking and isolation steps were repeated at least three times, and the 16S rRNA gene sequences and four housekeeping genes (*pheS, radA, gap*, and *ast*) were confirmed using Sanger sequencing performed using a BigDye Terminator v3.1 Cycle Sequencing Kit on Applied Biosystems 3130 genetic analyzer (ThermoFisher Scientific, Waltham, Mssachusetts, USA). The isolate was designated as the host strain, *A. pernix* YK1-12-2013.

### Virus culture and isolation

Approximately 5 g sample collected in 2014 were inoculated into fresh 1,000 mL JXTm medium in 2000 mL Erlenmeyer flasks sealed with silicone plug and incubated as described above. After 3 days, samples were centrifuged at 5,000 × g for 30 min in 500-ml Nalgene™ PPCO centrifuge bottles (ThermoFisher Scientific, Waltham, Mssachusetts, USA) at 4°C, and NaCl and polyethylene glycol 6000 were added to the supernatant (final concentration: 1 M and 10%, respectively). After incubation at 4°C overnight, viral particles were collected by centrifugation at 12,000 × g for 30 min at 4°C and suspended in 10 m: virus storage buffer (20 mM Tris-acetate at pH 7.0 containing 3.0% sodium chloride) and designated as the environmental virus fraction.

To screen viruses infecting A. *pernix* YK1-12-2013, we inoculated the 10 mL environmental virus fraction with 1,000 mL exponentially growing *A. pernix* YK1-12-2013 culture in 2000 mL Erlenmeyer flasks. As a control and mock treatment, we added virus suspension buffer and JXTm medium to the culture medium. Viral fractions were prepared as described above after further growth for approximately four days. Virus propagation was verified using transmission electron microscopy (TEM) as described below. For further purification, 10 mL chloroform was added. After vigorous vortexing, the suspension was centrifuged (7,500 ×g, 4°C, 20 min), and the aqueous layer was layered on a CsCl step-gradient (1.15, 1.25, and 1.40 g ml^-1^) and purified by CsCl step-gradient ultracentrifugation at 107,000×g for 60 min at 15°C using a Beckman Coulter Optima L-80 ultracentrifuge, SW41Ti rotor (Beckman Coulter Inc, Brea, California, USA). The concentrated viruses were collected using a 26-gauge needle and dialyzed in 500 mL SM buffer (50 mM Tris-HCl, 100 mM NaCl, 10 mM MgSO_4_ · 7H_2_O, 0.01% gelatin) at 4°C overnight.

### TEM analysis

For negative staining, 5 µL samples were applied to carbon-coated copper grids (Nisshin EM, Tokyo, Japan) stained with 2% uranyl acetate for 5–10 s. They were visualized using an H-7650 (Hitachi, Tokyo, Japan) at 80 kV at magnifications of 10,000× to 40,000×.

### DNA extraction, sequencing, and host and viral genome assembly

Extraction of cellular DNA and viral DNA was conducted using Wizard® Genomic DNA Purification Kit (Promega, Madison, Wisconsin, USA) and QIAamp MinElute viral DNA extraction kit (QIAGEN, Venlo, Netherland), respectively, following the manufacturer’s protocol. The sequencing library was prepared using a Nextrera XT library preparation kit (Illumina, San Diego, California, USA), and genomes were sequenced on the Illumina MiSeq v2 system with paired-end reads. The sequence of the viral genome was technically replicated (*n* = 2, defined as VL1 and VL2). We eliminated the reads with a Phred score below Q30 for 90% of the bases using the FASTQ Quality Filter of FASTX-Toolkit (http://hannonlab.cshl.edu/fastx_toolkit/). The reads were assembled using the Velvet Optimiser ver. 2. 2. 5 (http://bioinformatics.net.au/software.velvetoptimiser.shtml) (27).

To obtain the complete viral genome, VL1 and VL2 were assembled. We searched for homologous sequences between the VL1-contigs and VL2-contigs using the BLASTN program at the National Center for Biotechnology Information (NCBI) and assembled the contigs sharing significant homologous sequences using MEGA7 (28). If the length of the homologous sequence was less than 200 bp, the integrity of the assembly was confirmed using PCR. Finally, the gaps between the contigs were bridged using Sanger sequencing of the PCR products.

### Genome analysis

Open reading frames (ORFs) were predicted using the Microbial Genome Annotation Pipeline (MiGAP) (29) following manual curation. To determine the homology of ORFs to known proteins, we searched for non-redundant database protein sequences in the NCBI database using BLASTP (as of March 2021, *E* = 1e-5). Searches for conserved protein domains were performed using HMMscan (30) against Pfam (31) (as of March 2021. *E* = 1e-5). Membrane-spanning regions were predicted using the TMHMM ver. 2.0 (http://www.cbs.dtu.dk/services/TMHMM/) (32). Circular genome maps and GC skew calculations were performed using DNAPlotter (33). Genome alignments with related viruses based on genome-wide sequence similarities computed by tBLASTx were drawn using ViPTree web server version 1.9 (http://www.genome.jp/viptree/) (34).

To determine the AGV1-integration site in the host genome, we searched for homologous sequences between the virus genome and the *A. pernix* YK1-12-2013 draft genome using BLASTN. To ensure high sensitivity, sequence reads of the host were mapped onto the viral genome using Bowtie2 (35). The CRISPR elements and spacers were identified using CRISPRFinder (36) with manual validation. The virus genome was searched against the spacers registered in the CRISPRs database (as of March 2021) and the identified spacers of *A. pernix* YK1-12-2013 using BLASTN optimized for somewhat similar sequences.

### Protein analysis of virus

Purified virions were incubated at 95°C for 20 min with the buffer solution without the reducing reagent (6x) and subjected to SDS-PAGE (12% polyacrylamide gel; 80 × 90 × 1.0 mm; 200V; Nacalai Tesque, Kyoto, Japan) using ATTO AE-6450 mini PAGE system (ATTO, Tokyo, Japan). Proteins were transferred onto an Immobilon P membrane (Millipore, Burlington, Massachusetts, USA) using TRANS-BLOT SD SEMI DRY TRANSFER CELL (Bio-Rad, Hercules, California, USA) at 180 mA for 80 min and stained with CBB Stain One (Nacalai Tesque, Kyoto, Japan). A protein ladder (Nacalai Tesque, Kyoto, Japan) with molecular masses ranging from 10 to 250 kDa was used. The major protein band was excised using a scalpel, and the N-terminal amino acid sequence was determined by Edman degradation (APRO Science, Tokushima, Japan).

### Growth kinetics analysis of virus

We monitored the cellular (16S rDNA) and viral (integrase) genome copy numbers using qPCR. *A. pernix* YK1-12-2013 (1,000 mL culture) was grown in 2,000 mL Erlenmeyer flasks sealed with silicone plugs at 90°C without shaking. After 24 h incubation, 10 mL viral fraction or virus storage buffer (control; 20 mM Tris-acetate at pH 7.0 containing 3.0% sodium chloride) was added to *A. pernix* YK1-12-2013 cultures, and 10 mL was sampled every 24 h. For each sample, *A. penix* YK1-12-2013 was collected as described above. Virions were collected by ultracentrifugation at 25,000 rpm for 90 min at 4°C using a Beckman Coulter Optima L-80 ultracentrifuge, SW41Ti rotor. Cellular DNA and viral DNA were extracted from the cell-free supernatant by using kits as described above, and qPCR was performed for quantification. For the archaeal 16S rRNA gene, primer sets 931f-m1100r were used (37). The primers for the viral integrase gene were designed to target the 226-bp internal region of the integrase gene (Table 1). Standards used to determine the gene copy numbers of the 16S rRNA and integrase genes were prepared using the genomic DNA from *A. pernix* YK1-12-2013 and the purified virus, respectively. qPCR was performed using SYBR Premix Ex Taq II (Tli RNaseH Plus) (TaKaRa Bio, Shiga, Japan) on a TaKaRa PCR thermal cycler Dice® real-time system single and software. Thermal cycler Dice Real Time System Single (ver. 4.02 B for TP850) was used for the calculation of Ct values, generation of standard curves, and analysis of dissociation curves. Each PCR mixture (20 μL per tube) contained primers and SYBR® Green I solution of the recommended concentration according to the manufacturer’s guidelines. The PCR protocol for 16S rRNA genes was as follows: 60 s at 95°C for initial denaturation, 45 cycles of 5 s at 95°C, 10 s at 64°C, and 30 s at 72°C. The PCR protocol for integrase genes was as follows: 60 s at 95°C for initial denaturation, 45 cycles of 10 s at 95°C, 15 s at 61°C, and 40 s at 72°C. Dissociation curves were generated by gradually increasing the temperature from 60°C to 95°C after the PCR cycle.

**Table 1.**
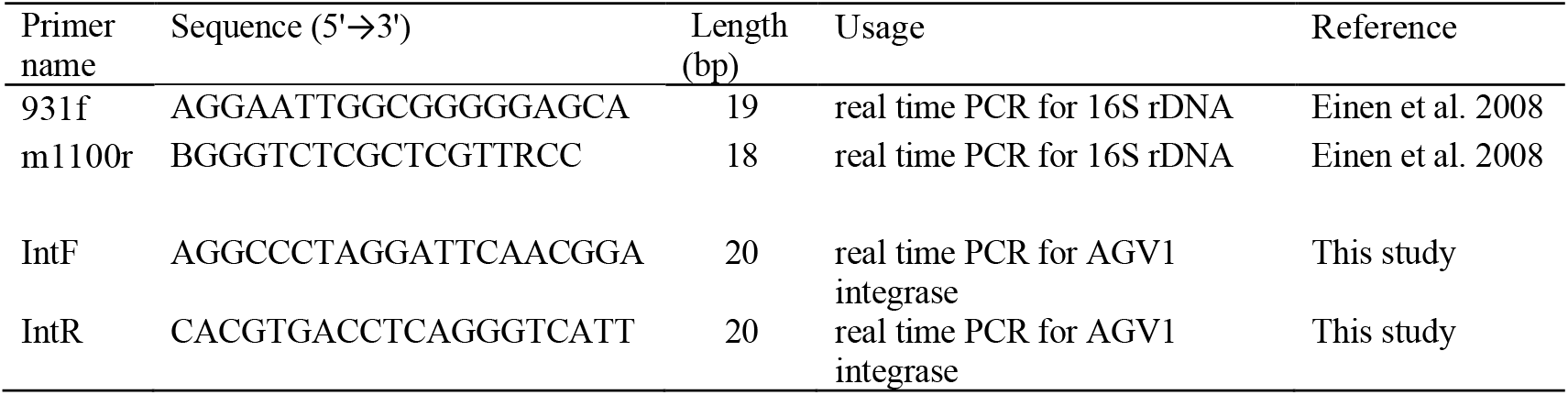
Primers used in this study.

### Growth analysis of host

The effect of viral infection on *A. pernix* YK1-12-2013 growth was confirmed using culture-turbidity measurements. Briefly, 100 µL AGV1 fraction, virus storage buffer, or JXTm was added to exponentially growing host cells in 5 mL culture in 18 × 180 mm hermetically sealed screw-cap test tubes. Every 8 h, 100 µL of culture was collected, and OD_600_ was measured using Ultrospec 3100 pro (GE Healthcare, Chicago, Ilinois, USA).

### Virus induction assay

Cells were grown in 5 mL culture in 18 ml screw-capped test tubes at 90°C without shaking for 24 h. To determine the factor inducing AGV1 replication, we performed stress treatments on *A. pernix* YK1-12-2013 cultures. First, pH was changed from 7.0 to 6.0 by adding 3M acetate. Second, exponentially growing host cells were transferred to shallow plastic dishes and subjected to UV irradiation at 254 nm for 30 s using a BioRad GS gene linker (BioRad, Hercules, California, USA). Third, we shifted the cultivation temperature from 90°C to 80°C. Fourth, 100 µL 5 M NaCl was added. Lastly, 100 µL 0.25 M EDTA (pH 7.0) was added. For each culture, cells were collected 48 h post-treatment, and cellular DNA was extracted and used as a PCR template to confirm AGV1 induction.

### Host range analysis

Host analysis was conducted using the following strains: *A. pernix* K1 and *A. camini* SY1 obtained from the Deutsche Sammlung von Mikroorganismen und Zellkulturen (DSMZ, Braunschweig, Germany) as DSMZ 11879 and DSMZ 16960, respectively, and three *A. pernix* laboratory strains isolated from various hydrothermal fields: OH2, TB7 (26), and FT1-29-2014 (from Shimogamo hot spring, Shizuoka, Japan).

Each strain was inoculated with 100 µL AGV1 fraction growing in 5 mL medium JXTm (defined as culture A) for 2 d. Then, 100 µL culture A was transferred to 5 mL fresh medium and incubated for another 2d twice (defined as cultures B and C). Genomic DNA was extracted from cultures A, B, and C per strain. Using these DNA templates, qPCR was performed using AGV1-specific primers to confirm infection. PCR using 16S rRNA-specific primers was used as the control.

### Data availability

The AGV1 genome and *A. pernix* YK1-12-2013 draft genome sequences have been deposited in the DNA Data Bank of Japan (DDBJ) database under the accession numbers LC208019 and BDMD01000001-BDMD010000144, respectively.

## RESULTS & DISCUSSION

### Isolation of novel temperate virus

An enrichment culture was established from samples collected from a hydrothermal field in Kagoshima, Japan, wherein *Aeropyrum* species and their infectious viruses have been previously isolated (26). From the TEM images, we observed spindle-shaped, pleomorphic, and linear virus-like particles in a viral fraction prepared from the enrichment culture (data not shown). To isolate the infectious viruses of *A. pernix*, the viral fraction was inoculated with *A. pernix* YK1-12-2013. After 4 days, host growth retardation was detected, and spherical particles approximately 60 ± 2 nm in diameter were observed in viral fractions from the cultures inoculated with virus fraction. And the same particles were also observed in viral fraction from culture added with virus storage buffer (Figs. 1). No virus-like particles were observed in the supernatant of *A. pernix* YK1-12-2013 cell culture without viral fraction or virus storage buffer. These observations indicate that the spherical viral particles are derived from a temperate virus (or viruses) within the *A. pernix* YK1-12-2013 strain.

**Fig. 1.**
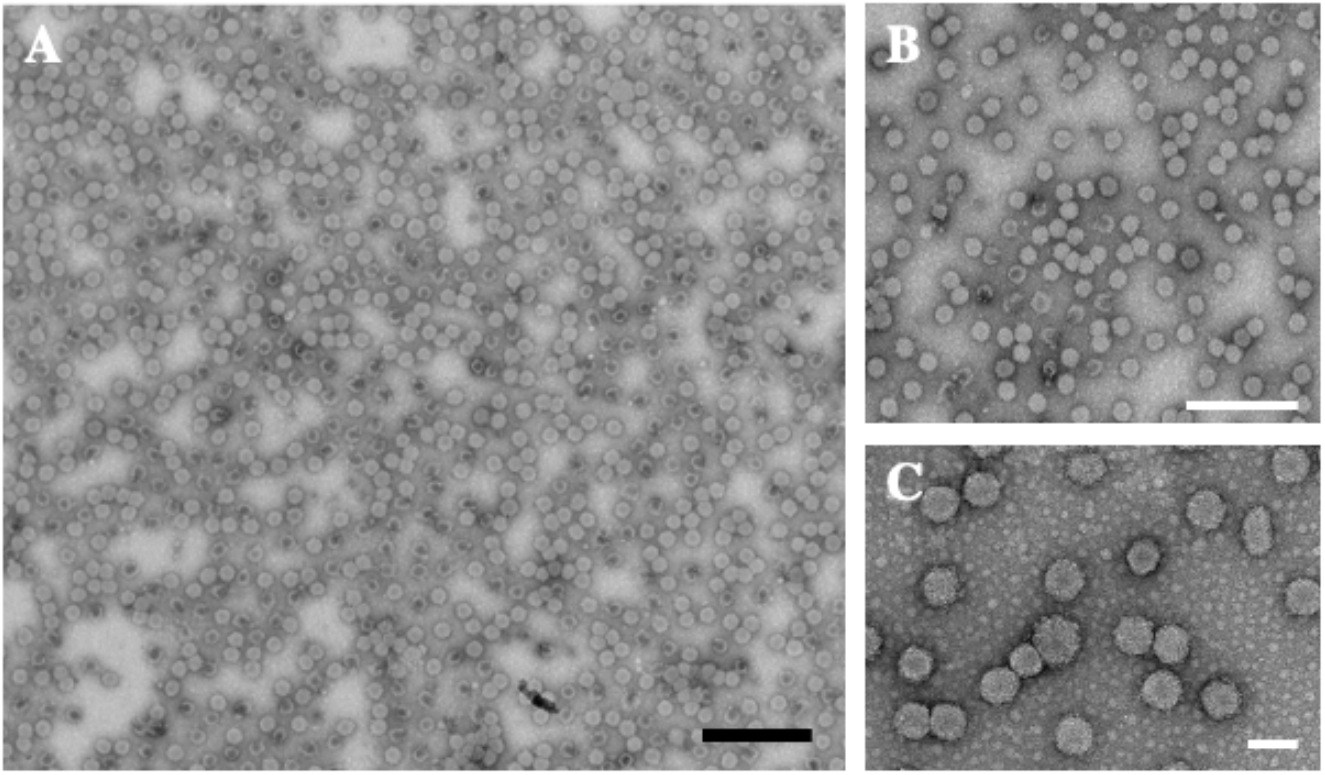
Morphology of AGV1 virions. Representative transmission electron micrographs of virus particles derived from *A. pernix* YK1-12-2013 cultures grown with buffer A. Cells are negatively stained with 1% uranyl acetate. Scale bars: 500 nm (A, B) and 100 nm (C).

The spherical virions are morphologically similar to those belonging to the *Globuloviridae* family, i.e., *Pyrobaculum* spherical virus 1 (38) and *Thermoproteus tenax* spherical virus 1 (39). Neither surface structures nor tails were observed. As there has been no reports to date on a spherical virus infecting *Aeropyrum* and we successfully obtained a single circular genome from these spherical virions later, we described these isolates as a novel virus species infecting *Aeropyrum* named *Aeropyrum* globular virus 1 (AGV1).

### AGV1 genomic features of and host genomic interactions

A single circular dsDNA genome containing 18,222 bp (Fig. 2A and Table 2) was assembled from spherically shaped particles. We numbered the nucleotides in the genome sequence beginning from the start codon of the first ORF following the replication origin. The ORFs are correspondingly numbered and the following number refers to the number of amino acids in the predicted protein (e.g. ORF1_309). The GC content (55.8%) was similar to those of host chromosomes (50.7–56.7%) (25) and the reported dsDNA viruses infecting *A. pernix* (52.7–56.5%) (15, 23)) but higher than those of other spherical viruses PSV (48%) (38) and TTSV (49.8%) (39). The AGV1 genome encoded 34 predicted ORFs starting from the ATG, GTG, and TTG codons (Table 3). Of these, 27 (79.4%) were present in one DNA strand, and only 7 were present on the other. The GC skew analysis revealed that the origin of replication is most likely located in the intergenic region between ORF 1_309 and ORF 34_215. We detected an inverted repeat, 5′-CACCCTGCTCTACTATAGTCTAT – N_93_– ATAGACTCTATAGTAGAGCCAGGGTG –3′ in this region. The *oriC* sites on the *A. pernix* K1 genome contain crenarchaeal origin recognition boxes (ORBs) which are the binding sites for Orc1/Cdc6 proteins and ori-specific uncharacterized motifs (UCMs), an important signature motif in the center of origin in *Aeropyrum* (40). We did not find any ORBs or UCMs around the putative replication origin on the AGV1 genome, suggesting that AGV1 cannot replicate autonomously. Of the predicted ORFs, 10 (29.4%) showed significant similarity to the genes in the nr database (Table 3). SDS-PAGE revealed a single major protein band of approximately 28.5 kDa from the protein fraction extracted from purified AGV1 virions (Fig. 2B), 5 N-terminal amino acid sequences of which completely matched with the internal amino acid sequences of a protein encoded by AGV1 ORF1_309. According to the HMMscan (30) analysis, 12 (35.3%) ORFs showed similarity to the protein domains registered in the Pfam database (31). We collectively assigned functions to these 13 AGV1 gene products (Fig. 2A, Table3).

**Table 2.**
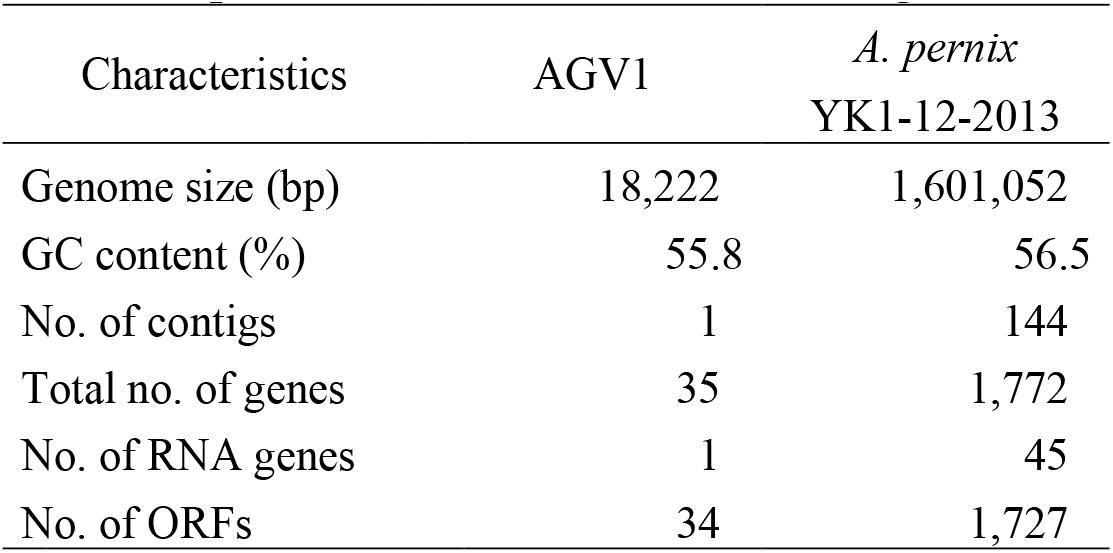
Sequence status of the host and AGV1 genomes.

| Characteristics | AGV1 | *A. pernix*<br>YK1-12-2013 |
| --- | --- | --- |
| Genome size (bp) | 18,222 | 1,601,052 |
| GC content (%) | 55.8 | 56.5 |
| No. of contigs | 1 | 144 |
| Total no. of genes | 35 | 1,772 |
| No. of RNA genes | 1 | 45 |
| No. of ORFs | 34 | 1,727 |

**Table 3.** Summary of the predicted ORFs in the AGV1 genome.

|  | position | Length (aa) | Function | HMM predictions | BLASTP Hit | Identity (%) | E-value |
| --- | --- | --- | --- | --- | --- | --- | --- |
| ORF1_309 | 1..930 | 309 | Structural protein* |  |  |  |  |
| ORF2_61 | 936..1121 | 61 |  |  |  |  |  |
| ORF3_233 | 1227..1928 | 233 |  |  |  |  |  |
| ORF4_262 | 1930..2718 | 262 | serine protease | Trypsin-like peptidase domain<br>(aa 57-197 hit to PF13365.4; 3.6e-21) | *Pyrodictium* sp<br>(ref\|HID40799.1) | 64/182(35%) | 1.00E-23 |
| ORF5_70 | 2708..2920 | 70 |  |  |  |  |  |
| ORF6_154 | 2913..3377 | 154 |  |  |  |  |  |
| ORF7_317 | 3361..4314 | 317 |  |  |  |  |  |
| ORF8_354 | 4329..5393 | 354 |  |  |  |  |  |
| ORF9_64 | 5390..5584 | 64 |  |  |  |  |  |
| ORF10_110 | 5600..5932 | 110 |  |  |  |  |  |
| ORF11_119 | 5925..6284 | 119 |  |  |  |  |  |
| ORF12_80 | 6427..6669 | 80 |  |  |  |  |  |
| ORF13_229 | 6927..7616 | 229 |  |  |  |  |  |
| ORF14_163 | 7784..8275 | 163 |  |  |  |  |  |
| ORF15_146 | 8272..8712 | 146 | DNA binding (RHH) | Ribbon-helix-helix protein, copG family<br>(aa 111-147 hit to PF1402.21; 4.6e-07) |  |  |  |
| ORF16_133 | 8957..9358 | 133 |  |  |  |  |  |
| ORF17_88 | 9388..9654 | 88 |  |  |  |  |  |
| ORF18_111 | 9660..9995 | 111 |  | Uncharacterized protein conserved in bacteria<br>(aa 3-109 hit to PF09995.9; 3.2e-06) | *Aeropyrum pernix* ovoid virus 1<br>(ref\|WP_010865925.1\|) | 70/111(63%) | 1.00E-42 |
| ORF19_114 | 10003..10347 | 114 | DNA polymerase | DNA polymerase alpha subunit p180 N terminal<br>(aa13-69 hit to PF12255.8; 1.8e-06) |  |  |  |
| ORF20_141 | 10355..10780 | 141 | pillin | Archaeal Type IV pilin, N-terminal (aa 4-93 hit to PF07790.11; 1.0e-08) | *Thermogladius calderae* 1633 (ref\|AFK51634.1\|) | 59/151(39%) | 9.00E-19 |
| ORF21_191 | 10777..11352 | 191 | nuclease | Viral/Archaeal nuclease (aa 22-175 hit to PF12187.8; 3.3e-13) | *Pyrodictium delaneyi* (ref\|WP_082419610.1\|) | 75/180(42%) | 2.00E-28 |
| ORF22_70 | 11363..11575 | 70 | DNA binding (RHH) | Ribbon-helix-helix protein, copG family (aa 21-58 hit to PF01402.21; 5.0e-10) | *Aeropyrum pernix* spindle-shaped virus 1 (ref\|YP009177778.1\|) | 45/69(65%) | 5.00E-24 |
| ORF23_123 | 11578..11949 | 123 |  |  |  |  |  |
| ORF24_108 | 11959..12285 | 108 |  | HemN C-terminal domain (aa 9-77 hit to PF06969.16; 1.9e-06) |  |  |  |
| ORF25_82 | 12248..12496 | 82 |  | TIR domain (aa 4-60 hit to PF13676.6; 1.4e-06) | *Aeropyrum pernix* (ref\|WP_131159643.1\|) | 51/86(59%) | 3.00E-26 |
| ORF26_339 | 12493..13512 | 339 | integrase | Archaeal phage integrase (aa 180-333 hit to PF16795.5; 2.4e-40) | *Sulfolobus* spindle-shaped virus 1 (ref\|NP_039778.1\|) | 117/333(35%) | 1.00E-44 |
| ORF27_64 | 13509..13703 | 64 |  |  |  |  |  |
| ORF28_309 | complement(13710..14639) | 309 | ATP binding protein | P-loop containing region of AAA domain (aa 28-65 hit to PF13555.6; 1.7e-06) | *Pyrodictium* sp.(ref\|HID41144.1\|) | 142/249(57%) | 9.00E-93 |
| ORF29_107 | complement(14642..14965) | 107 |  |  |  |  |  |
| ORF30_111 | complement(14967..15302) | 111 |  |  |  |  |  |
| ORF31_258 | complement(15377..16153) | 258 | Glutamyl-tRNA<sup>Glu</sup> reductase | Glutamyl-tRNA<sup>Glu</sup> reductase, dimerisation domain (aa 64-143 hit to PF00745.20; 1.3e-06) | *Aeropyrum pernix* ovoid virus (WP_010865937.1) | 189/258(73%) | 2.00E-133 |
| ORF32_80 | complement(16160..16402) | 80 |  |  |  |  |  |
| ORF33_330 | complement(16407..17399) | 330 |  |  | *Thermoprotei archaeon* (ref\|RLE66374.1\|) | 42/106(40%) | 7.00E-08 |
| ORF34_215 | complement(17421..18068) | 215 |  |  |  |  |  |

**Fig. 2.**
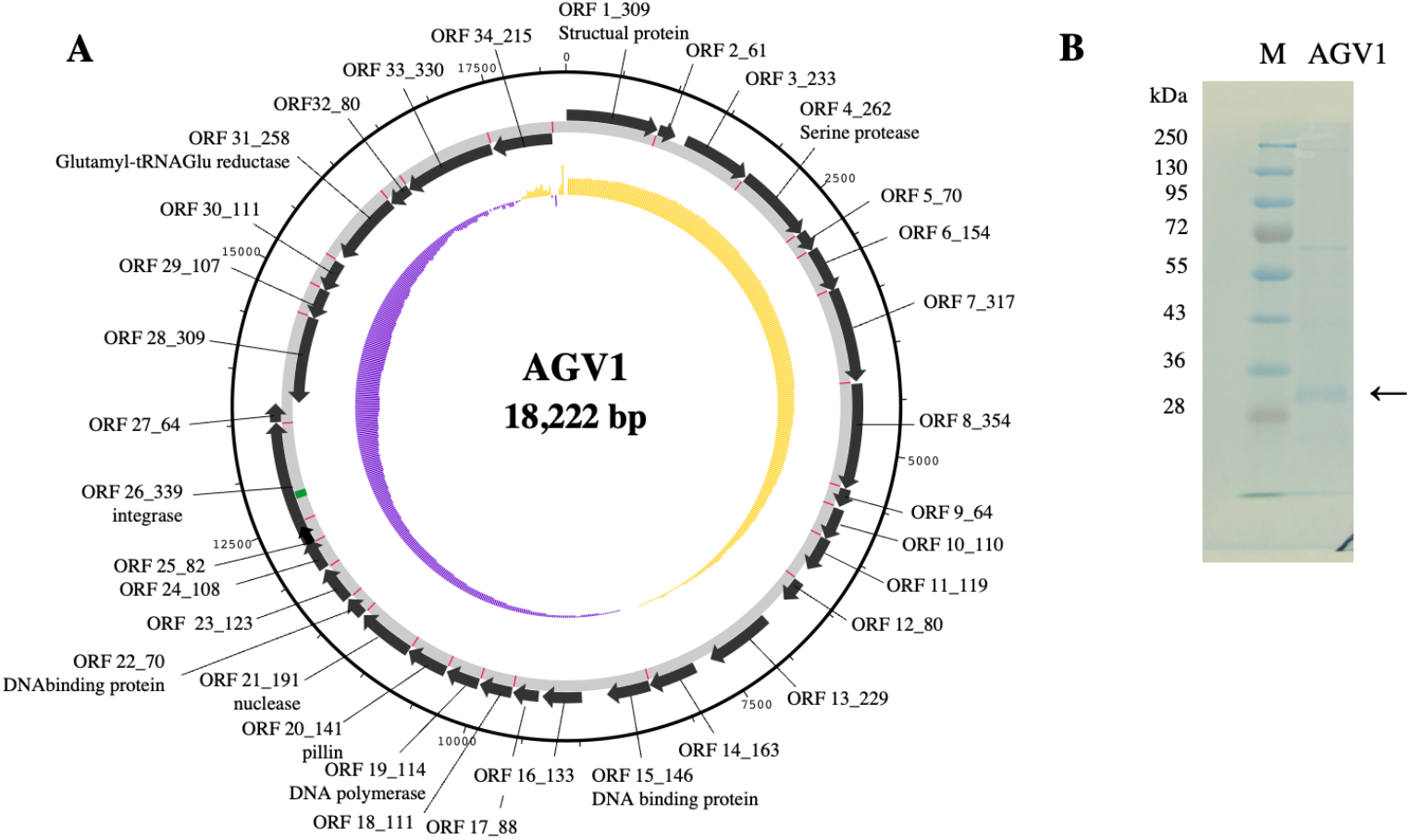
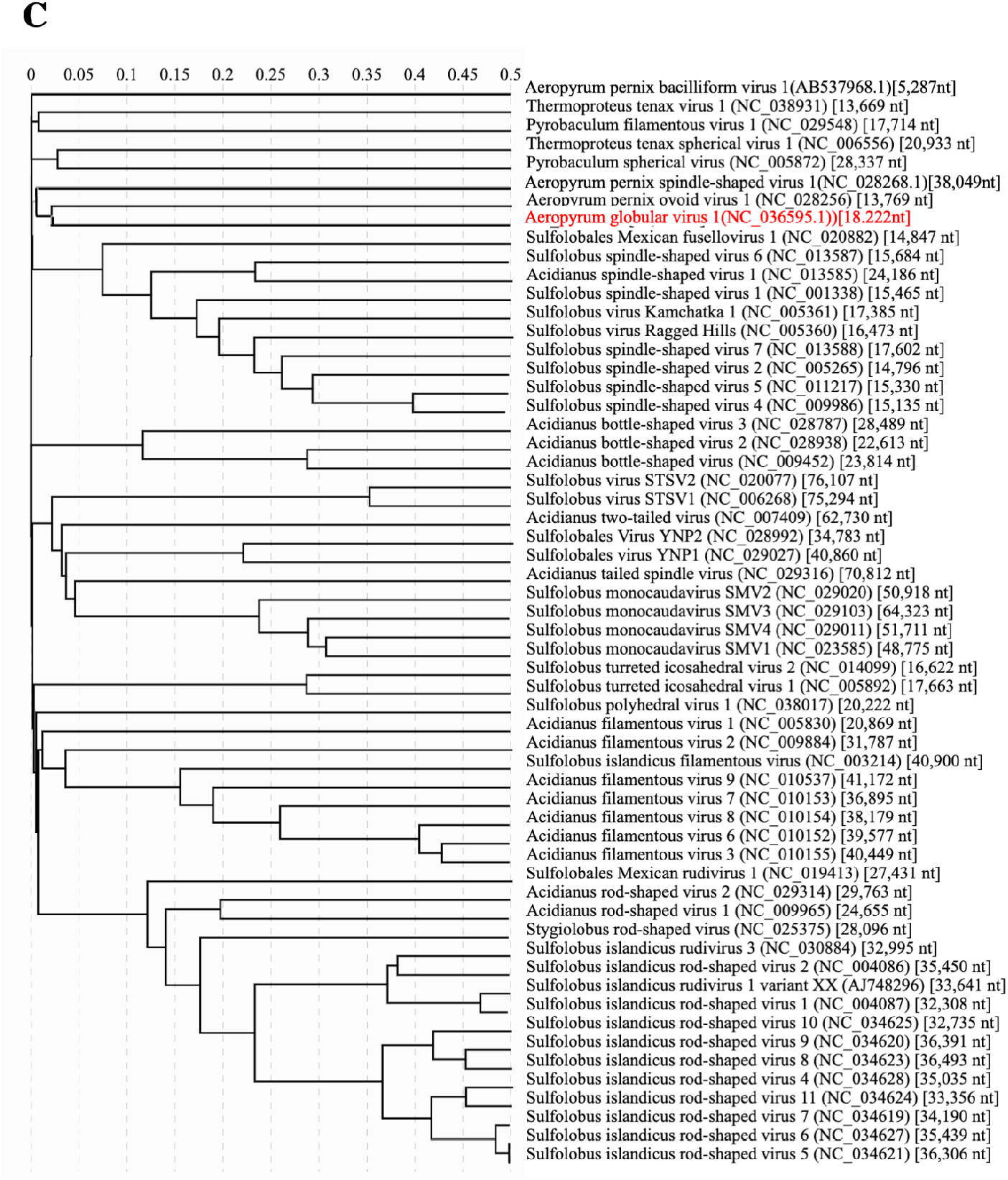
Genomic information of AGV1. (A) Genome map of AGV1. Outer ring shows the genomic contents: black arrows, ORFs with strand orientation; red, ribosome-binding site; and green, tRNA^Glu^ gene. Inner ring shows the GC skew (window size: 9000 and step size: 25): purple, negative GC skew; yellow, positive GC skew. Replication origin is located at the intergenic region between ORF1_309 and ORF34_215. (B) Structural protein of AGV1. Representative SDS-PAGE gel shows the proteins of the virions stained with CBB stain one. Molecular mass marker (M) are shown. The arrow indicates the viral major protein identified. (C) Phylogenetic tree of the genetic relationship among crencarchaeal viruses based on genome-wide sequence similarities computed by tBLASTx. The tree was generated by ViPTree. SG values are shown in top row.

PSV1 and TTSV1, the reported members of *Globuloviridae*, share 15 genes including two structural proteins (39). Although AGV1 virions are morphologically similar to PSV1 and TTSV1, the structural protein of AGV1 showed no similarity with those of PSV1 and TTSV1. Moreover, no homologs were found on the AGV1 genome. The five AGV1 genes showed significant similarities with those of reported crenarchaeal temperate viruses. Particularly, three of the five genes are similar to that in Fuselloviruses, namely integrase (ORF 26_339) in the SSV-like region on *Sulfolobus islandicus* M.16.4, nuclease (ORF 21_191) in SSV2, and putative transcriptional regulator (ORF 22_70) in APSV1. Integrase genes found in archaeal extrachromosomal elements are classified into two types: SSV1-type and pNOB8-type (40). AGV1 integrase is an SSV1-type integrase, a kind of tyrosine recombinase catalyzing the site-specific integration and excision of the viral genome (40). HMMS revealed that the functional domain was conserved in the integrase gene. We found that the tRNA^Glu^ gene was inserted in the AGV1 integrase gene sequence, suggesting that tRNA^Glu^ is the integration site of the integrase. Furthermore, ORF22_70 products encoded a putative DNA-binding protein containing the ribbon-helix-helix (RHH) domain common to the CopG family. This domain was also predicted for the ORF15_146 product. The RHH domain protein is one of the most conserved proteins in archaeal viruses (41) and recognizes target sequences in the DNA by forming dimers and inserting them into the groove (42). The RHH domain was also found in F55, which tightly regulates the SSV1 lysogenic state (20) and is involved in controlling plasmid copy numbers (42).

Four ORFs of AGV1 were homologs of *Aeropyrum-*encoded genes, three of which (ORF18_111, ORF22_70, and ORF31_258) are shared with the provirus on the *A. pernix* K1 genome. AGV1 shares a hypothetical protein (ORF18_111) and putative glutamyl tRNA^Glu^ reductase (ORF31_258) with APOV1.

Five ORFs showed sequence similarity to those of other hyperthermophilic crenarchaeota, including trypsine-like peptidase (ORF4_262), nuclease (ORF21_191), and ATP-binding protein(ORF28_309) in *Pyrodictium* sp.; pillin(ORF 20_141) in *Thermogladius calderae*; and hypothetical protein (ORF33_330) in *Thermoprotei archaeon*. Trypsine-like peptidase is also encoded by the ssDNA virus ACV (24) and might be involved in the release or invasion cycle of these viruses.

ORF28_309 exhibited a P-loop containing region of the AAA domain, which is conserved among the ATP- or GTP-binding proteins and is encoded by many archaeal viruses (43). Typically, viral P-loop proteins are nucleic-acid-stimulated ATPases that are involved in viral replication, transcription, or packaging and are related to bacterial DnaA (43). We found that 13 ORFs, including structural proteins and pilli, exhibit one to four transmembrane motifs predicted by the TMHMM 2.0 program.

Finally, we compared the AGV1 genome with those of related crenarchaeal viruses on a genome-wide scale using ViPTree (34). AGV1 shared some genes with APOV1 and *Fuselloviridae* members, but most of its genomic content was unique, indicating the novelty of AGV1. The genome architecture of AGV1 showed mosaicism, which may reflect the genetic exchange among other viruses and organisms. Genome mosaicism of viruses has been reported between bacterial phages and archaeal head-tail viruses. To date, there have been no reports on viruses infecting crenarchaeota (9, 44).

To predict the AGV1 integration site on the host genome, we performed a draft genome sequence analysis of the host. The draft assemblies of the *A. pernix* YK1-12-2013 genome yielded 144 contigs with an average GC content of 56.5% (Table 2). The draft genome was approximately 1.6 Mbp long, and a total of 1,727 ORFs were identified. However, we could not detect the AGV1 integration site in the host genome or any sequence reads of the host mapped to the AGV1 genome, including tRNA^Glu^. That is to say, the read coverage of the AGV1 genome was lower than that of the host genome indicating that AGV1 was harbored by some host cells in culture. APOV1 and APSV1 harbor the SSV1-type integrase and are integrated into the chromosome of the host *A. pernix* K1 (15). In addition, *Fuselloviridae* members encode SSV1-type integrases and are integrated into the host chromosome (40). In contrast, *Bicaudaviridae* members, which also carry the SSV1-type integrase but integration of viral genome has not yet been reported except for ATV(13). Furthermore, a recent study revealed that the integrase gene is not essential for SSV1 (45); thus, genome integration is not always essential for infection. We obtained a pure host cell culture using previous methods (22, 26) and confirmed no redundancy of 16S rRNA genes and housekeeping genes by Sanger sequencing and draft genome sequencing (data not shown but the sequences obtained by Sanger sequencing completely matched those of the draft genome). Therefore, we presumed that AGV1 is not integrated into the host genome and maintained as a episome in the host cell and that initially, the host culture consisted of only AGV1-carrying cells, but AGV1-carrying cells were diluted upon host cell division as AGV! could not replicate autonomously due to the lack of the ORBs or UCMs.

### Virus-host relationship

To confirm the maintenance and replication of the AGV1 genome in host cells, we monitored the host/virus genome copy numbers during infection. AGV1 genome copy number was 5.6 to 35 × 10^3^ times lower than that of host genome, and replication did not occur without induction stimulus (Fig. 3). At 9 h post-treatment, AGV1 in the host cells rapidly increased. At 48 h post-treatment, its genome copy number reached approximately 3 to 5 × 10^8^ copies mL^-1^ culture and exceeded that of host genomes by 9.5 to 26-fold. The AGV1 genome copy number in the supernatant exhibited a similar increasing pattern and plateaued at approximately 3×10^7^ copies mL^-1^ culture at 72 h post-treatment. When it plateaued, AGV1 virions were observed in the cell free supernatant by TEM.

**Fig. 3.**
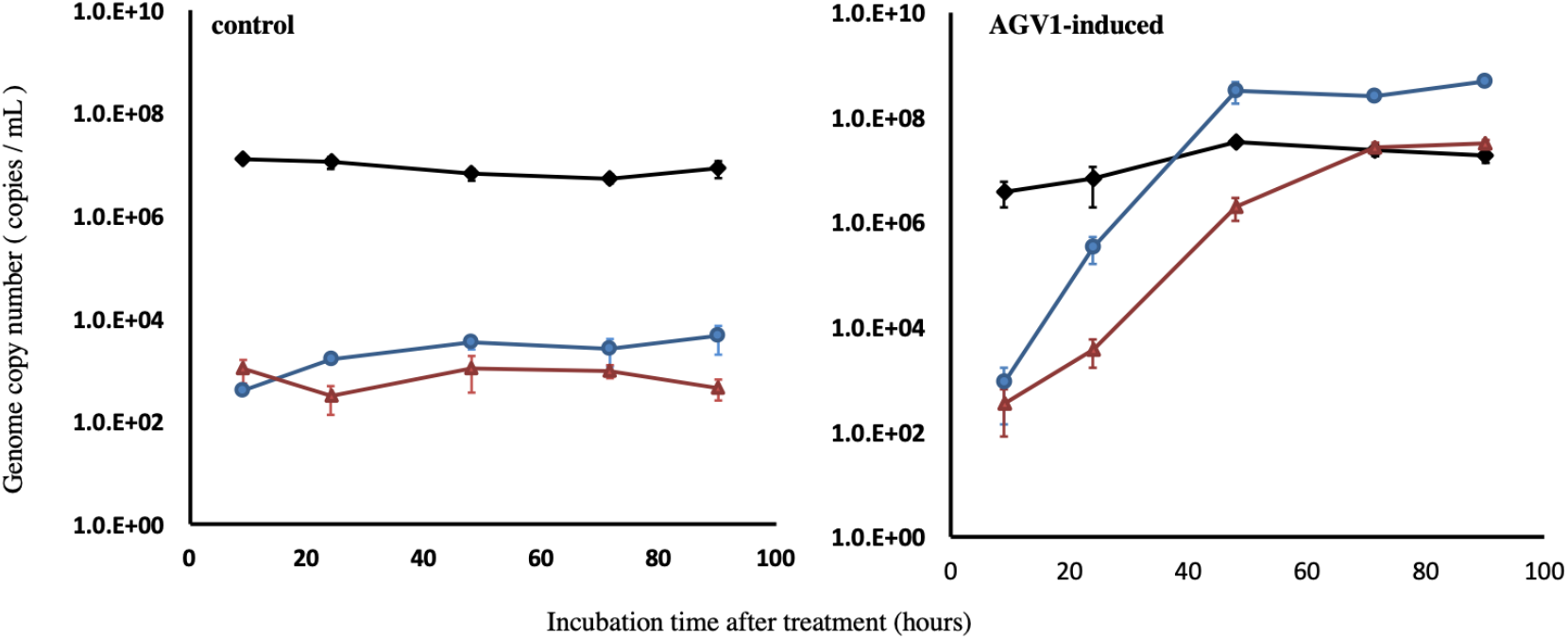
Time-course of AGV1 and host genome copy numbers. Native host *A. pernix* YK1-12-2013 was cultivated at 90°C, and samples were collected at the indicated time points. The host genome copy numbers (black diamond) and the virus genome copy numbers in the cell pellet (blue circle) and supernatant (red triangle) from 10 mL culture of *A. pernix* YK1-12-2013 were measured using quantitative PCR analysis of the DNA extracted from the host cells and released virions, respectively. Error bars represent the mean ± SE of triplicates.

To clarify whether AGV1 infection affects host growth, we monitored host growth with or without AGV1 and under AGV1-induced conditions. Host growth was more significantly retarded following AGV1 inoculation than under AGV1-induced conditions. Growth retardation was observed immediately after AGV1 inoculation and 12 h post-induction, at which AGV1 propagation in the host cell started (Fig. 4). Infection with AGV1 did not lead to host cell lysis as evidenced by a lack of decrease in OD_600_ and the absence of cell debris in the culture of infected cells.

**Fig. 4.**
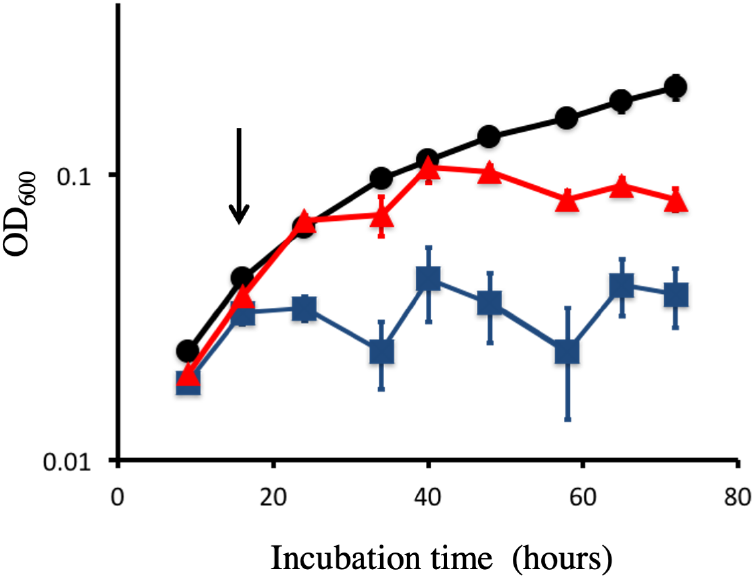
Effect of AGV1 on the growth of *A. pernix* YK1-12-2013 strain. Black circles, red triangles, and blue squares indicate the control, tris-acetate buffer-added culture, and AGV1 fraction-added culture. The AGV1 fraction or tris-acetate buffer was added at the time indicated by the arrow. Error bars represent the mean ± SE of six replicates.

### *Aeropyrum*–AGV1 interaction via the CRISPR/Cas system

CRISPR is a microbial adaptive immune system that cleaves foreign genetic elements (46). CRISPR allows cells to specifically recognize the target sequences using spacers acquired from the proto-spacer (small DNA segment of invaders) and inhibits viral infection, such as RNA interference manner (47). To investigate the AGV1–host interaction via CRISPR, we identified the CRISPR loci on the host genome and spacers originating from AGV1. *A. pernix* YK1-12-2013 carried two CRISPR loci (Apyk_1 and Apyk_2) carrying 39 and 57 repeat-spacer units, respectively.

We detected *cas*2 and *cas*4 genes adjacent to Apyk_2. *cas2* and *cas4* genes are important for insertion of spacers into CRISPR arrays. We also detected other *cas* genes involved in spacer acquisition (*cas1*), target binding (*csa2, cas5, cmr2*, and *cmr3*), and target cleavage (*cmr4, cmr6, and csx1*) in the draft genome (46). Direct repeat sequences and leader sequences of Apyk_1 and Apyk_2 were matched to the repeat sequences of Ape_1 and Aca_2 (25), which are previously reported CRISPR loci of *A. pernix* K1 and *A. camini* SY1, respectively. We found 8 protospacers on AGV1 (Table 4), 3 of which matched those on the CRISPR loci on the *A. pernix* YK1-12-2013 genome, whereas the remaining five matched those of *A. pernix* K1. The sequence of the single spacer (Apyk_2_45) perfectly matched that of the protospacer. Other protospacers had at least one nucleotide mutation.

**Table 4.**
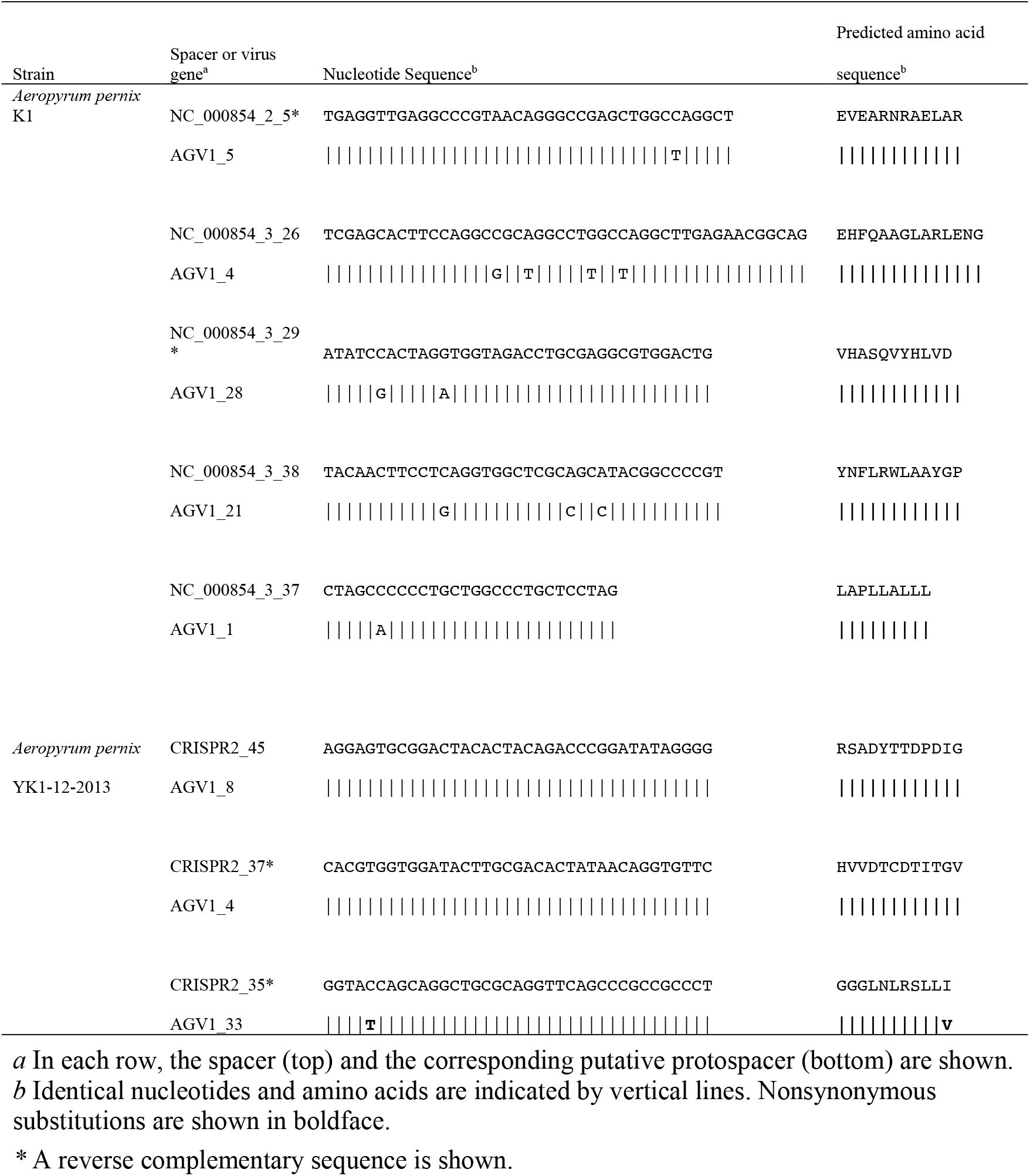
Comparison of spacer sequences to identify putative protospacers.

Next, the nucleotide mutations in the protospacers were examined to determine whether they caused synonymous substitutions. We found that most nucleotide mutations caused synonymous substitutions, except for Apyk_2_35. Apyk_2_35 contains a single amino acid substitution of isoleucine to valine; however, this residue was conserved, suggesting an arms race between AGV1 and the host. Our results suggest that CRISPR protects *A. pernix* YK1-12-2013 from AGV1 infection.

Recent transcriptomic analysis of *S. solfataricus* infected with two closely related temperate viruses, SSV1 and SSV2, indicates that CRISPR interacts with the temperate virus and regulates its copy number (19). SSV2 showed a steep increase in copy number (from to 1–3 to 25–50 copies per cell) during host growth (48) and elicited a strong host response, including CRISPR (19). On the other hand, SSV1 maintained its copy number at a low level without any induction stimulus (18) and did not activate the host defense system (19). Furthermore, viruses with destructive effects on the host induce a stronger host defense response as evidenced by the high numbers of spacer matches to lytic virus ATV found in the CRISPR loci of *S. solfataricus* P2 compared with carrier-state viruses (e.g., fuselloviruses infecting the strain) (9). Thus, we assumed that the CRISPR/Cas immunity of *A. pernix* YK1-12-2013 represses AGV1 propagation because high concentrations of AGV1 might have damaging effects on host growth. Alternatively, AGV1 might be in the cryptic phase during the optimal growth phase of the host to escape from the host defense. However, AGV1 can propagate steeply upon certain stress stimuli, such as addition of the virus storage buffer. The detailed mechanism of AGV1 life cycle remains to be elucidated.

### Factors associated with AGV1 induction

To identify the AGV1 induction stimulus, we examined AGV1 propagation under various stress conditions. PCR products of AGV1 were obtained in cells growing under stress conditions stimulated, namely suboptimal growth pH, suboptimal growth temperature, and UV irradiation (Fig. 5) besides tris addition. Salinity shift and addition of chelating agent didn’t induce AGV1 propagation. AGV1 genome copy number was approximately measured as 1 × 10^3^ copies mL^-1^ cultures under suboptimal growth pH, suboptimal growth temperature and UV irradiation. The addition of tris-acetate buffer induced approximately 1 × 10^8^ copies mL^-1^ and was the most effective stimulus to induce AGV1 propagation.

**Fig. 5.**
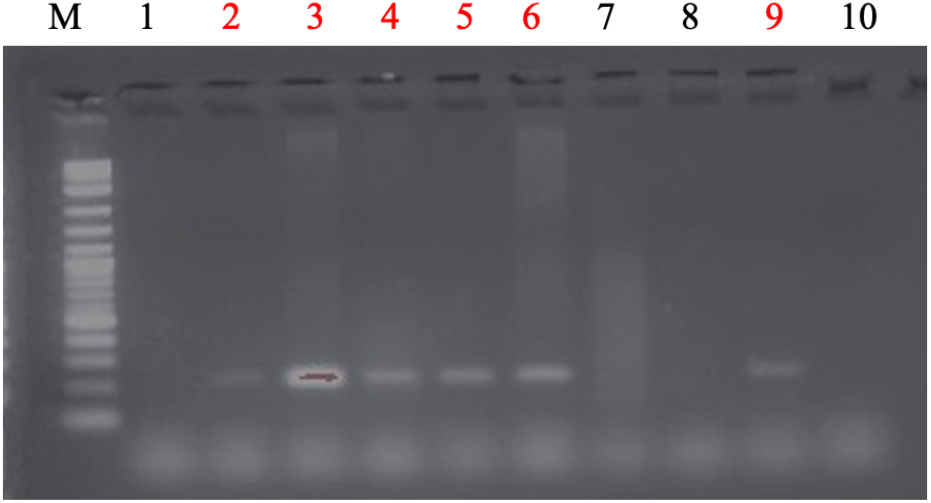
AGV1 Induction assay. PCR analysis was performed using AGV1-specific primers to confirm the kind of stress induced by AGV1 propagation. M: 2 log ladder, 1-9: YK1-12-2013 genome extracted from cells under various stress treatments: 1: optimal, 2: addition of 20 mM tris-acetate buffer, 3: addition of 50 mM tris-acetate buffer, 4: addition of 100 mM tris-acetate buffer, 5: UV irradiation for 30 s, 6: suboptimal growth temperature, 7: addition of 20 mM EDTA (pH 7.0), 8: high salinity, 9: suboptimal growth pH, and 10: DDW (negative control). The lanes confirming product amplification are in red.

### Host range analysis

The host range of AGV1 was evaluated by inoculating AGV1 with exponentially growing cells of five *Aeropyrum* strains isolated from various hydrothermal fields in Japan. We did not detect the AGV1 genome in all cultures of OH2, TB7, and FT1-29-2014 strains. In *A. pernix* K1, we detected the AGV1 genome at a low level (4.8 to 7.0 × 10^2^ copies mL^- 1^) in cultures A and C. In *A. camini* SY1, we detected the AGV1 genome at 1.0 × 10^6^ copies mL^-1^ in culture B and at 5.5 × 10^6^ copies mL^-1^ in culture C. 16S rRNA gene was detected in all cultures at approximately 10^5^ copies mL^-1^.

Thus, *A. pernix* K1 and *A. camini* SY1 could be infected by AGV1. We continued to subculture both AGV1-infected strains and after 10-time subculturing, we performed qPCR targeting AGV1 integrase. However, we could not detect the AGV1 genome in cultures with or without the virus storage buffer. Similarly, the *Sulfolobus neozealandicus* droplet-shaped virus belonging to *Guttaviridae* was found in an unstable carrier state (16). In contrast, we did not obtain any AGV1-free strain of the native host, *A. pernix* YK1-12-2013, despite long-term subculturing and effort to curation by plating and limiting dilution. Elucidation of the detailed mechanism should be explored in future studies.

Overall, we successfully isolated a novel spherical temperate virus AGV1 that infects the hyperthermophilic archaeon *Aeropyrum*. AGV1 was morphologically similar to *Globuloviridae* viruses but doesn’t share any genes with them. According to the genetic information, AGV1 seemed to be a temperate virus but AGV1 couldn’t neither integrate into the host genome nor replicate autonomously. On the other hand, AGV1 could propagate with inducing stimulus in spite of threat of the host CRISPR/Cas system. Although an “unstable carrier state” of AGV1 seemed to be disadvantageous for the survival, it might be a reasonable way to adapt to the host defense system and the harsh environment of the host habitat. We hypothesize AGV1 could hides from host defense system by unstable carrier state without inducing stimulus and propagate steeply on detecting stressful condition for its host. These results and hypothesis shed new light on virus-host interaction under extreme environment.

## ACKNOWLEDGMENTS

Computational analysis was performed at the Super Computer System, Institute for Chemical Research, Kyoto University. We thank the Center for Anatomical, Pathological, and Forensic Medical Researches (Graduate School of Medicine, Kyoto University) for their technical assistance with transmission electron microscopy. We also thank Takashi Daifuku and Shin Fujiwara for their assistance in host isolation experiments and genomic analysis. The authors declare that they have no conflicts of interest. This work was supported by a Grant-in-Aid for Challenging Exploratory Research (grant number 26640112) from the Japan Society for the Promotion of Science. The funders had no role in study design, data collection and interpretation, or the decision to submit the work for publication. We would like to thank Editage (www.editage.com) for English language editing.

